# Development of attenuated coxsackievirus B3 vectored intranasal pre-emptive pan-coronavirus vaccine

**DOI:** 10.1101/2023.11.22.568225

**Authors:** Huixiong Deng, Xuanting He, Yanlei Li, Haoyang Wang, Shenmiao Wang, Liming Gu, Hengyao Zhang, Jiacheng Zhu, Rui Li, Gefei Wang

**Affiliations:** Guangdong Provincial Key Laboratory of Infectious Diseases and Molecular Immunopathology, Shantou University Medical College, Shantou, Guangdong 515041, China

**Author notes:** These authors contributed equally (Huixiong Deng, Xuanting He, Yanlei Li). Correspondence: Dr. GeFei Wang; Dr. Rui Li.

**Keywords:** Pan-Coronavirus, Intranasal immunization, multivalent, epitope, mucosal vaccine

## Abstract

SARS-CoV-2 has the ability to evade immunity, resulting in breakthrough infections even after vaccination. Similarly, zoonotic coronaviruses pose a risk of spillover to humans. There is an urgent need to develop a pre-emptive pan-coronavirus vaccine that can induce systemic and mucosal immunity. Here, we employed a combination of immune-informatics approaches to identify shared immunodominant linear B- and T-cell epitopes from SARS-CoV-2 variants of concern (VOCs) and variants of interest (VOIs), as well as zoonotic coronaviruses. Epitope-guided vaccine were designed and the attenuated coxsackievirus B3 vectored intranasal vaccines rCVB3-EPI and rCVB3-RBD-trimer were constructed. The immunogenicity of these candidate vaccines was evaluated using Balb/c mice. The results demonstrated effective immune responses, including the production of SARS-CoV-2-specific IgG and sIgA antibodies, as well as T cell-mediated responses. However, further verification is required to assess cross-reactivity with various variants. Our intranasal pre-emptive pan-coronavirus vaccine design framework offers an appealing candidate for future vaccine development.

## Introduction

Currently available COVID-19 vaccines have shown high efficacy in preventing severe disease. However, they may not provide sufficient protection against infection or virus transmission over time. It is evident from the continued spread of SARS-CoV-2 in highly vaccinated populations, particularly with the emergence of new variants [1,2]. The ongoing transmission and replication of the virus allow for the evolution and emergence of immune-evasive variants, which can reduce the effectiveness of current vaccines in preventing infection and symptomatic disease [2–5]. On the other hand, the origin of deadly coronaviruses can be traced back to bats, and their transmission to humans occurs through various intermediate animal reservoirs. Given the genetic diversity of these coronaviruses and the risk of cross-species infection, there is a high possibility of future global COVID pandemics caused by spillover events from bat-derived coronaviruses. Such events pose significant threats to public health [6]. Therefore, it is essential to develop a safe and effective pre-emptive pan-coronavirus vaccine that targets a wide range of human and animal coronavirus strains. This vaccine should aim to reduce infection and transmission of zoonotic coronaviruses while maintaining or enhancing protection against symptomatic and severe disease. To achieve these goals, mucosal vaccination via the respiratory route is a logical approach to elicit robust mucosal immunity in the respiratory tract [7].

Building upon our previous work on a multivalent enterovirus subunit vaccine based on immunodominant epitopes [8], we employed similar approaches in this study. Specifically, we designed three all-in-one epitope-based pre-emptive pan-coronavirus vaccine candidates using highly conserved genome-wide human B- and T-cell epitopes from seven antigenic proteins of SARS-CoV-2. These candidates are expected to provide protection not only against COVID-19 but also against future zoonotic coronavirus outbreaks. Additionally, we utilized a genetically stable and temperature-sensitive attenuated live CVB3 vector, engineered with attenuated mutations (A78E) in the VP1 and (F237Y) in the VP3 structural protein [9]. In a mouse model, this viral vector demonstrated invasion of the mucosa and effectively induced both mucosal and systemic immune responses through nasal mucosal immunization. It also showed robust vertical maternal-infantile immune protection (data not published), serving as a platform for both delivery vesicles and mucosal immune stimulators. We constructed the candidate vaccine based on coxsackievirus B3 vectors and administered it via the nasal mucosal route to preliminarily evaluate its immunogenicity using a mouse model. This evaluation aimed to determine its efficacy in providing immune protection locally and systemically, thus laying the foundation for the development of a pre-emptive multi-epitope pan-coronavirus vaccine. This vaccine would be capable of preventing past, present, and unforeseen strains with epidemic or pandemic potential.

## Results

### The candidate immunodominant epitopes were highly conservative between SARS-CoV-2 VOCs and VOIs as well as zoonotic coronaviruses based on sequence characteristics

Based on computer-assisted vaccine design technology, we analyzed the sequence and structural features of coronaviruses to screen several immunodominant candidate epitopes, optimizing them, and selecting suitable delivery vectors and routes to develop a pan-coronavirus vaccine capable of inducing both systemic and mucosal immunity (Figure 1). To identify conserved immunodominant epitopes among zoonotic coronaviruses with a risk of human spillover, we conducted a screening of the evolutionary relationships and genetic similarities among 33 alpha- and beta-CoVs, which may pose a higher risk of spillover, including 8 of high-risk H-CoVs, 12 zoonotic high-risk bat-origin CoVs, and 13 CoVs from intermediate hosts during CoV zoonotic transmission, which including pangolin (7 CoVs), civet cat (1 CoV), camel (3 CoVs), murine (2 CoVs). The phylogenetic analysis of complete genomes revealed that the SARS-CoV-2 (Wuhan-Hu-1) strain is located in the inner part of the phylogenetic tree, clustering with BtCoV-RaTG13 and pangolin SL-CoVs, suggesting a consensus characteristic between SARS-CoV-2 and these coronavirus isolates. It is likely that the virus spilled over into humans after further mutations and/or recombination (Figure 2A). Sequence alignments showed genetic similarity ranging from 49% to 100% among the 33 CoVs. Pangolin-origin CoVs, Bat-SARS-like coronaviruses, SARS-CoV-Urbani, and Civet-SARS-CoV-007 exhibited high genetic similarity with SARS-CoV-2, while murine-origin CoVs showed maximum genetic distance and limited similarity to other CoVs (Figure 2B). Given the ongoing global pandemic caused by SARS-CoV-2, based on the evolutionary trajectory information of SARS-CoV-2 by phylogenetic tree and sequence alignment analysis, it is reasonable to consider SARS-CoV-2 as a reference isolate to design a pre-emptive pan-coronavirus vaccine capable of preventing current and unforeseen human outbreaks and deter future zoonotic events. To construct our dataset for SARS-CoV-2, we retrieved B-cell, Th-cell, and cytotoxic T-lymphocyte (CTL) immunodominant epitopes from the IEBD database, which have been defined for their immunological efficacy in several studies. These epitopes were mapped to the complete protein sequences of SARS-CoV-2. Using a combinatorial approach as described in the methods section, we selected candidate linear B-cell epitopes primarily derived from the Surface glycoprotein (10/12), Membrane glycoprotein (1/12), and Nucleocapsid (1/12). These epitopes have been shown to induce neutralizing antibodies that can block virus entry into target cells (Table 1). Notably, several candidate epitopes, such as B1 (467-513 aa), B9 (444-469 aa), B10 (439-454 aa), and B12 (402-424 aa), overlap with the receptor binding domain (RBD) region of the Spike protein, which binds to the ACE2 receptor. These epitopes can generate antibodies that inhibit conformational changes or the fusion process during viral entry. Sequence alignment and clustering analysis revealed that epitope B3 (Surface glycoprotein, 862-876 aa) was conserved in all 106 variants of VOC and VOI, while 11 B-cell epitopes exhibited a high degree of conservation among these variants, with most strains being 100% conserved and a few showing at least 75% conservancy (Table S1). For coronaviruses, 10 B-cell epitopes were found to be 100% conserved among two or more H-CoVs and SL-CoVs originating from pangolins and bats (GX-P4L, GX-P5E, GX-P5L, P2V, P1E, MP789, Bt-CoV-recombinant strain, Rs672, W1V1, and YNLF_31C). Additionally, 10 B-cell epitopes showed at least 70% conservation among two or more CoVs from previous outbreaks, such as SARS-CoV and SL-CoVs (SARS-CoV-Urbani, HCoV-OC43), as well as SL-CoVs originating from bats, pangolins, and camels (Civet007, BtCoV-HKU4, BtCoV-recombinant strain, Rs672, WIV1, Bat Hp-BetaCoV, Pipistrellus bat coronavirus HKU5 isolate YD13403, BtCoV-WIV16, BtCoV-RATG13, Rousettus BtCoV-GCCDC1, YNLF_31C, MERS-CoV, HCoV-EMC [MERS-related coronavirus strain]; GX-P1E, GX-P5E, GX-P4L, GX-P1E, GX-P5L, GX-P5E, GX-P2V, MP789) (Table S2). The presence of glycosylation is an important factor influencing virus immunogenicity and causing viral immune evasion. Upon screening for glycosylation regions, only two B-cell epitopes, B5 (Membrane glycoprotein, 1-15 aa) and B11 (Surface glycoprotein, 1186-1205 aa), were predicted to be N-glycosylated, while the remaining B-cell epitopes were found to be non-glycosylated (Supplementary data 1). These findings indicate that 12 candidate B-cell epitopes can provide coverage for 106 current SARS-CoV-2 epidemic variants and 33 CoVs with a high risk of human spillover. Regarding Th-cell epitopes, we selected 16 immunodominant CD4^+^ T-cell epitopes from our epitope dataset. These epitopes demonstrated strong binding to different MHC-II alleles and induced the release of specific cytokines, such as IFN-γ, TNF-α, IL-2, IL-5, and IL-4 (Table 2). Out of these candidates, 7 mapped to the Surface glycoprotein, followed by 6 in the Nucleocapsid protein. Conservation analysis showed that Th1, Th5, Th9, Th11, Th15, and Th16, which mapping to the Membrane glycoprotein (176-190 aa), Nucleocapsid protein (311-325 aa), Surface glycoprotein (866-880 aa), ORF8a (43-57 aa), Surface glycoprotein (896-910 aa), Nucleocapsid protein (107-121 aa), respectively, are 100% conserved among 106 SARS-CoV-2 variants (Table S3). Among the Th epitopes, 16 were 100% conserved among two or more H-CoVs and SL-CoVs. However, lower conservation was observed in the SL-CoV strain originating from civet and the MERS-CoV strain isolated from camels (Table S4). We identified 18 CD8^+^ T cell epitopes derived from the Surface glycoprotein, Nucleocapsid phosphoprotein, and open-reading-frames (ORF3a and ORF1ab) of SARS-CoV-2 (Table 3). The highest degree of similarity was observed among 106 SARS-CoV-2 strains, 8 strains of previous HCoVs, and 25 animal SL-CoV strains isolated from bats, civet cats, pangolins, mice, and camels. CTL1, CTL2, CTL3, CTL4, CTL5, CTL7, CTL9, CTL14, CTL15, and CTL16 exhibited 100% conservation across the 106 variants of VOC and VOI for SARS-CoV-2 (Table S5). A total of 18 CTL epitopes showed 100% conservation among two or more H-CoVs and SL-CoVs, including the four major "common cold" coronaviruses that caused previous outbreaks (HCoV-OC43, HCoV-229E, HCoV-HKU1, and HCoV-NL63), as well as the SL-CoVs isolated from bats and pangolins (Table S6).

**Figure 1.**
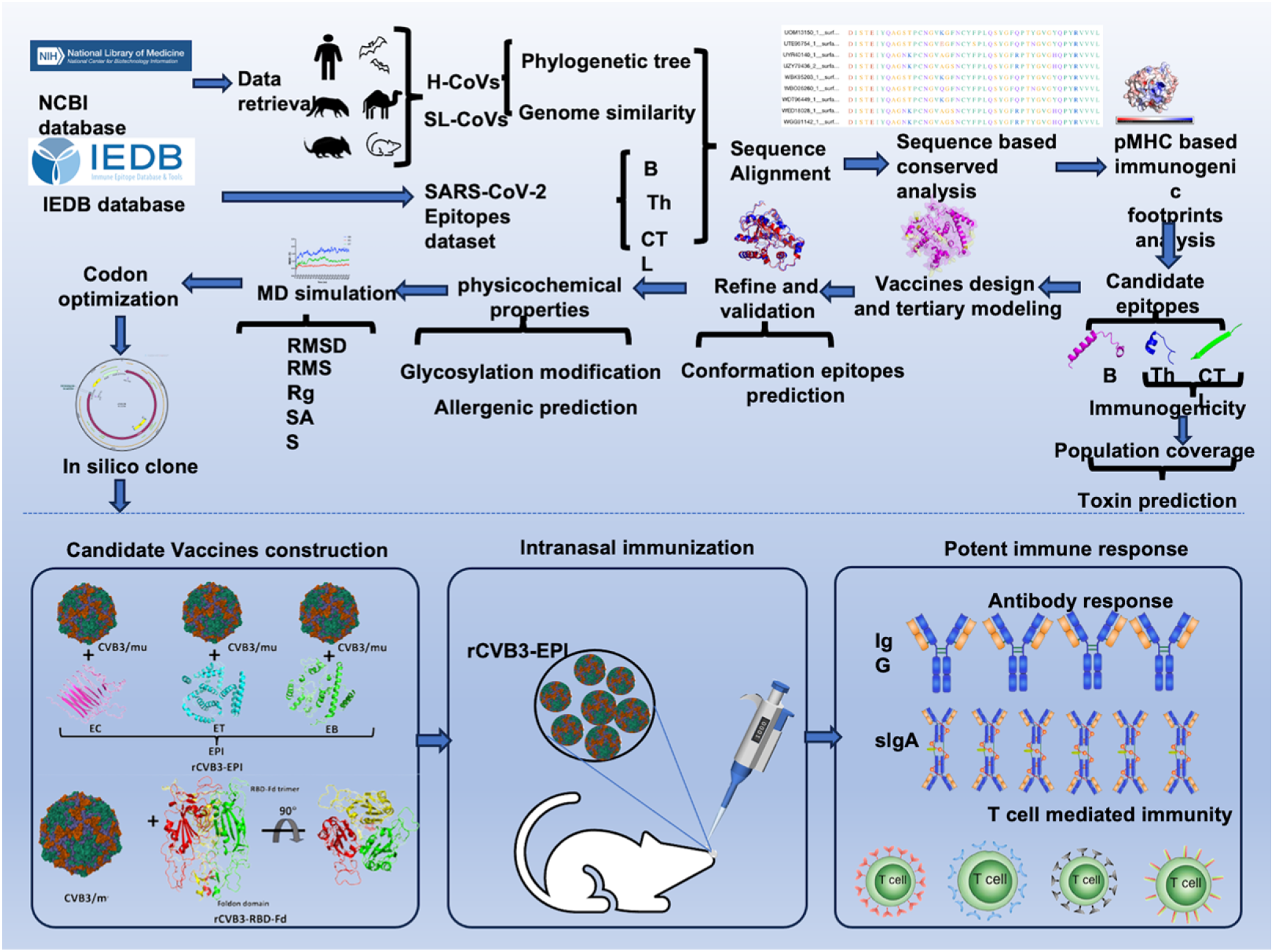
Schematic representation of the intranasal pre-emptive pan-coronavirus candidate vaccine based attenuated coxsackievirus B3 delivery vector designing process using immunodominant B-cell, CTL and Th epitopes.

**Figure 2.**
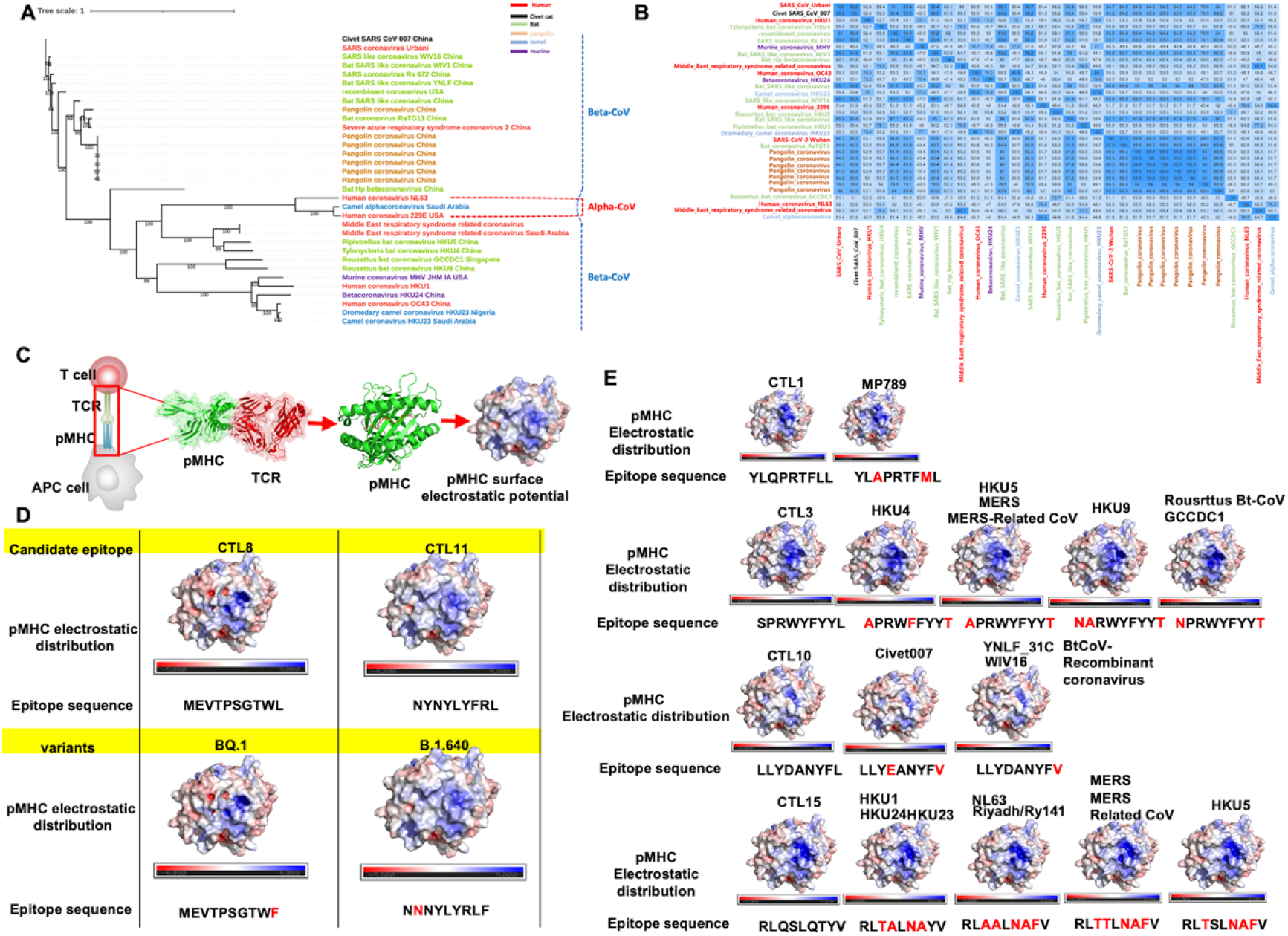
Evolutionary relationship and genetic similarity among 33 alpha- and beta-CoVs and the candidate CTL epitopes immunogenic footprints analysis. **(A)** Evolutionary comparison of genome sequences among Alpha- and beta-CoVs strains isolated from humans and animals’ host. A maximum-likelihood phylogenetic analysis was performed and generated 33 whole sequences between SARS-CoV-2 strains and seven isolates originating from human host (red, n=8), along with the animal’s SARS-like coronaviruses genome sequence (SL-CoVs) sequences origin from bats (green, n=12), pangolins (yellow, n=7), civet cats (black, n=1 ), camels (blue, n=3 ), and murine (purple, n=2). The included SARS-CoV/MERS-CoV strains are from previous outbreaks (origin from humans [HCoV-Urbani, MERS-CoV, HCoV-OC43, HCoV-NL63, HCoV-229E, HCoV-HKU1-genotype-B], bats [BtCoV-WIV16, BtCoV-WIV1, BtCoV-YNLF-31C, BtCoV-Rs672, BtCoV-recombinant strains], camel [Camelus dromedaries, (KT368891.1, MN514967.1, KF917527.1, NC_028752.1], and civet [Civet007, A022, B039]). **(B)** Homology of among 33 CoVs genome sequences originating from the human, bat, pangolin, civet cat and camel. Results indicate maximum genetic identity percentage among SARS-CoV-2 (Wuhan-Hu-1) strain and BtCoV-RaTG13 (97.2 %), pangolin coronavirus (92.3 %) and BtCoV-bat-SARS-like-coronavirus (91.1 %). **(C)** Schematic diagram of pMHC and TCR interaction. **(D)** Comparisons among pMHCs carrying candidate CTL epitopes and the correspondent variant sequences in SARS-CoV-2 VOCs and VOIs. Quality and reliable structural modelling of complexes predicted using Alphafold 2.3.2. DockQ score are up to 95. The electrostatic distributions surfaces from the candidate epitopes and correspondent variants are not different, even presenting amino acid changes in peptides suggesting a preserved TCR recognition. The variants BQ. 1and B.1.640 with their respective changes (marker Red) and correspondent epitope sequences. **(E)** Comparison of electrostatic surface distribution and topography of candidate CTL epitopes with coronaviruses putative epitopes. The variants with their respective changes (marker Red) and correspondent epitope sequences, electrostatic calculations are depicted as negative (red) and positive (blue) charges (-5 to +5).

**Table 1.**
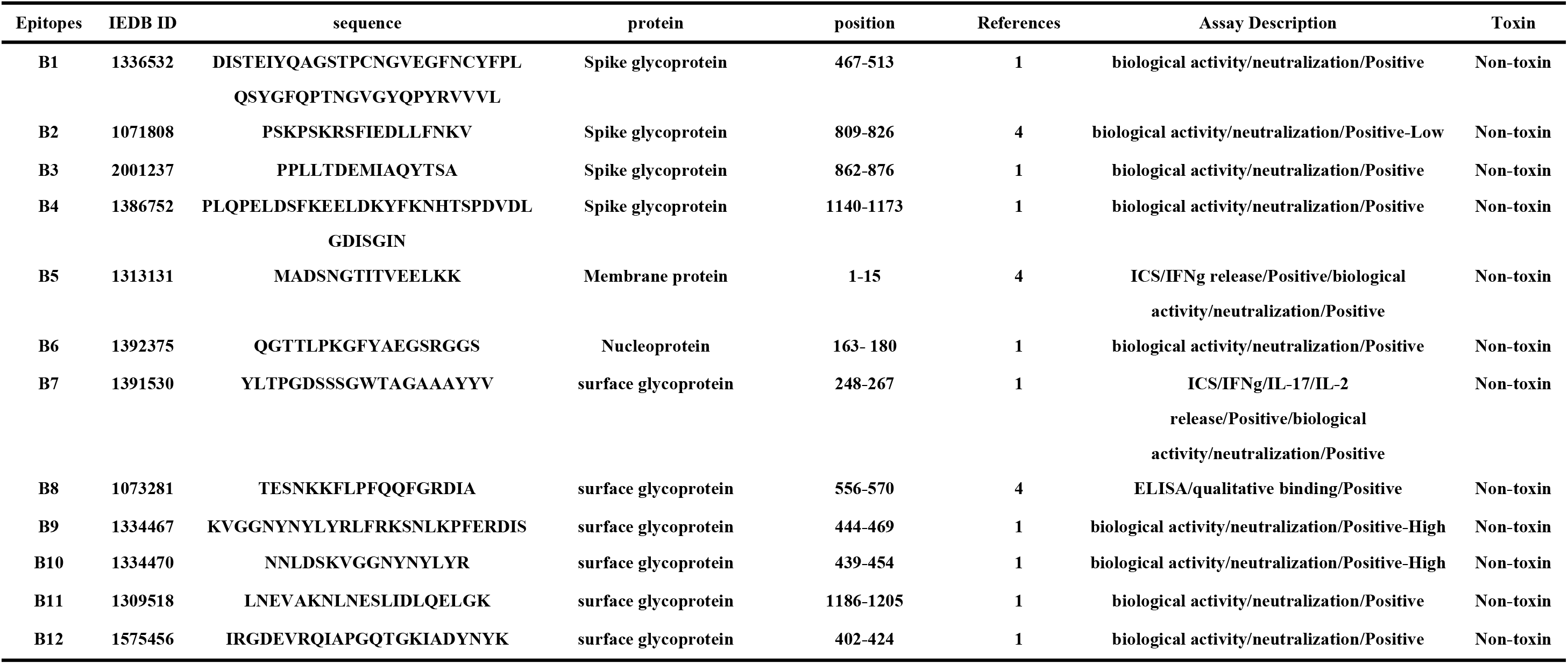
Linear B-cell candidate epitopes for the pre-emptive Pan-Coronavirus Vaccine as well as toxin prediction for each epitope.

**Table 2.**
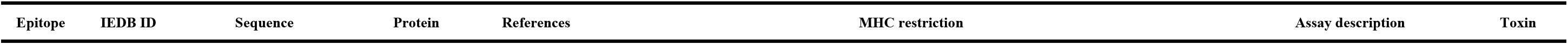
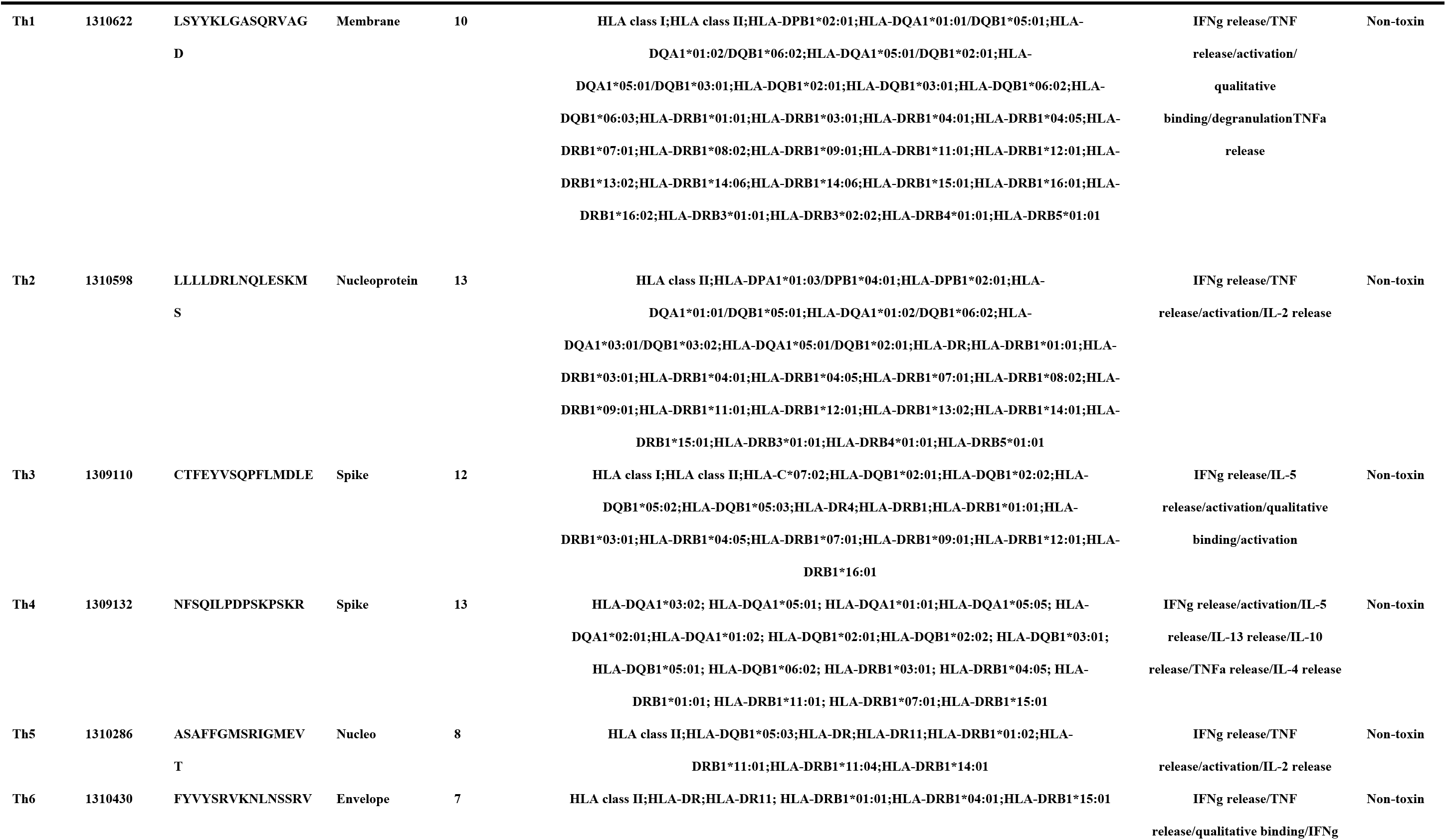
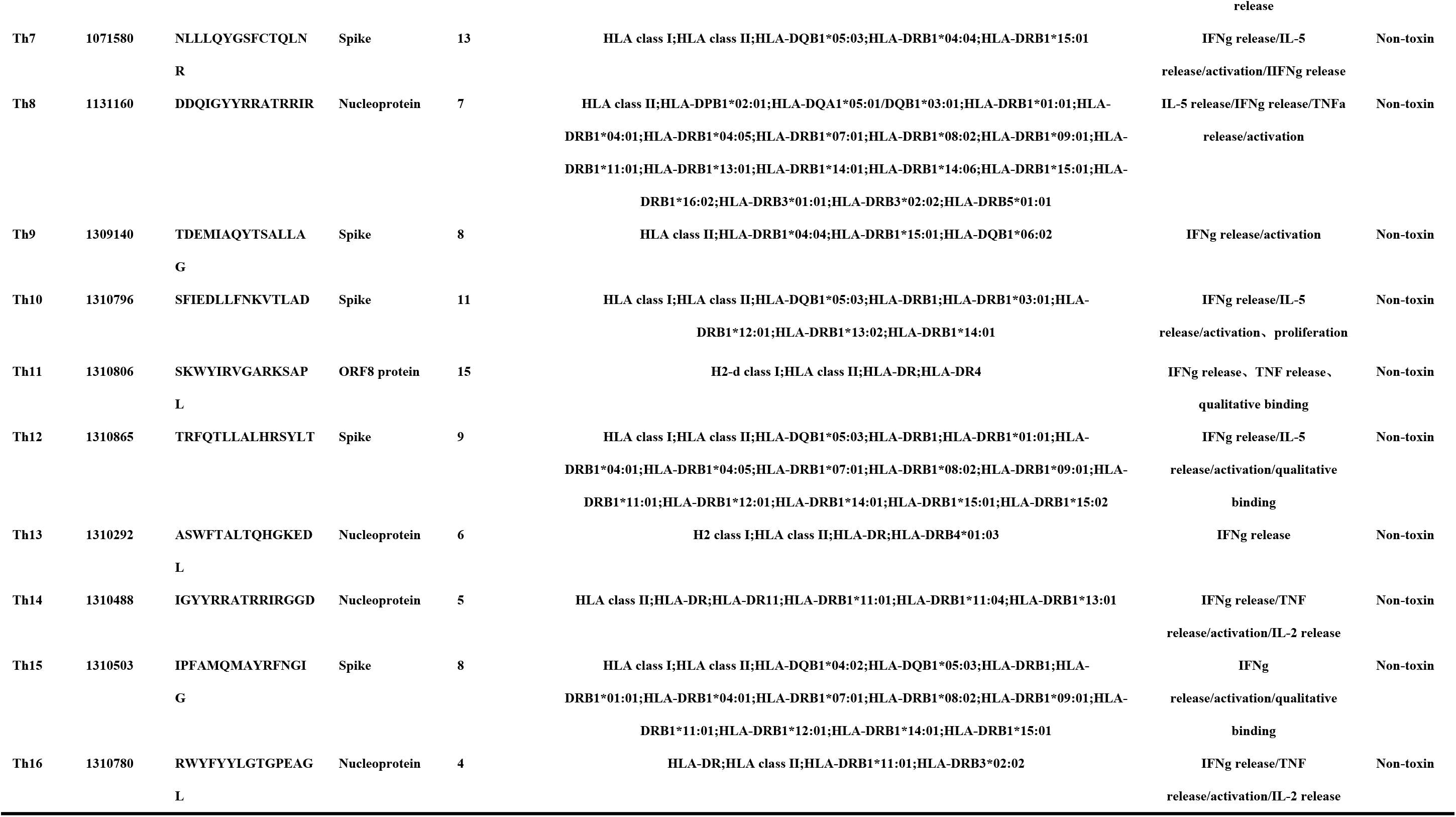
Helper T-lymphocyte specific epitope for the candidate pre-emptive Pan-Coronavirus Vaccine as well as toxin prediction for each epitope.

**Table 3.**
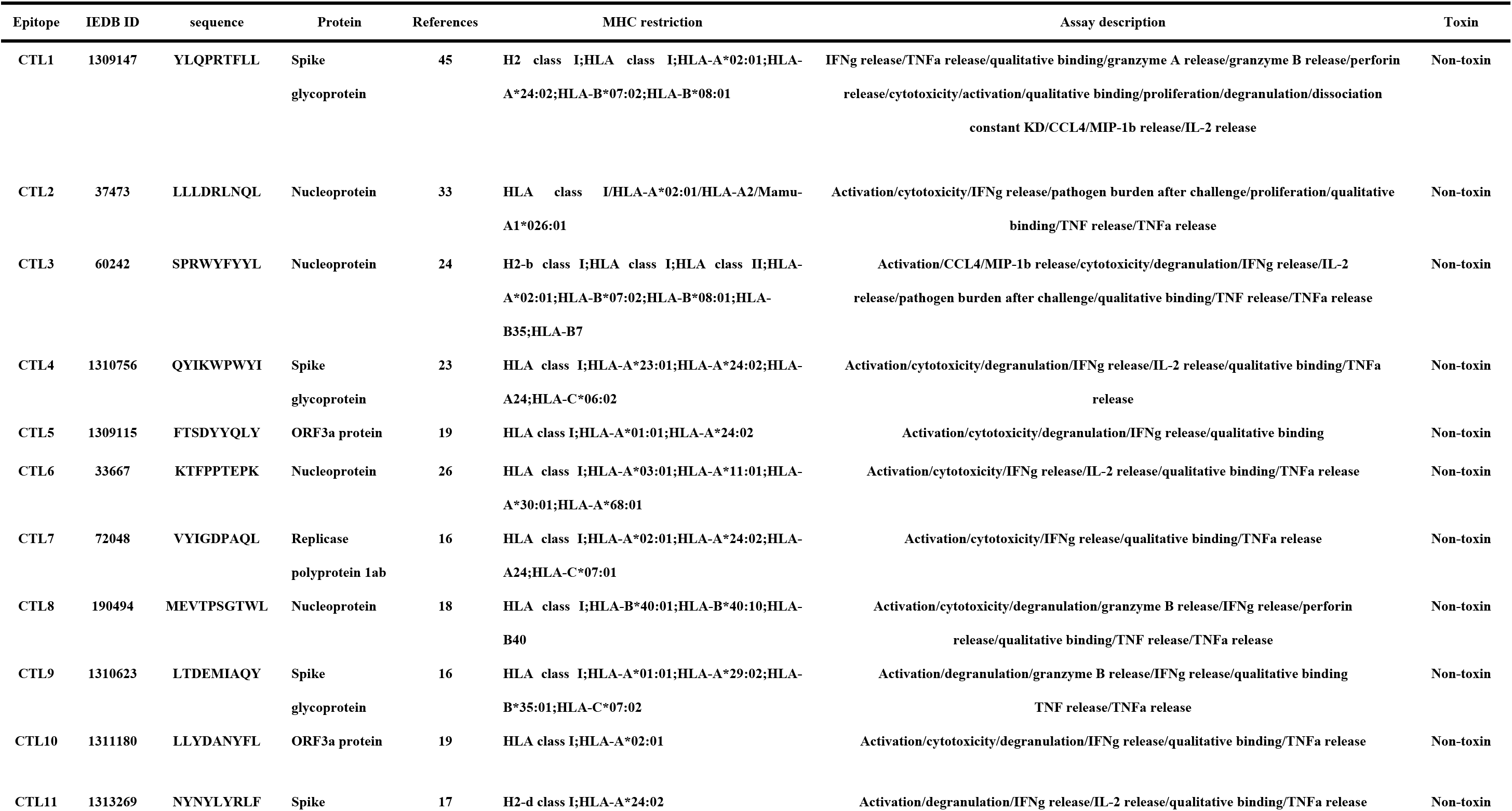
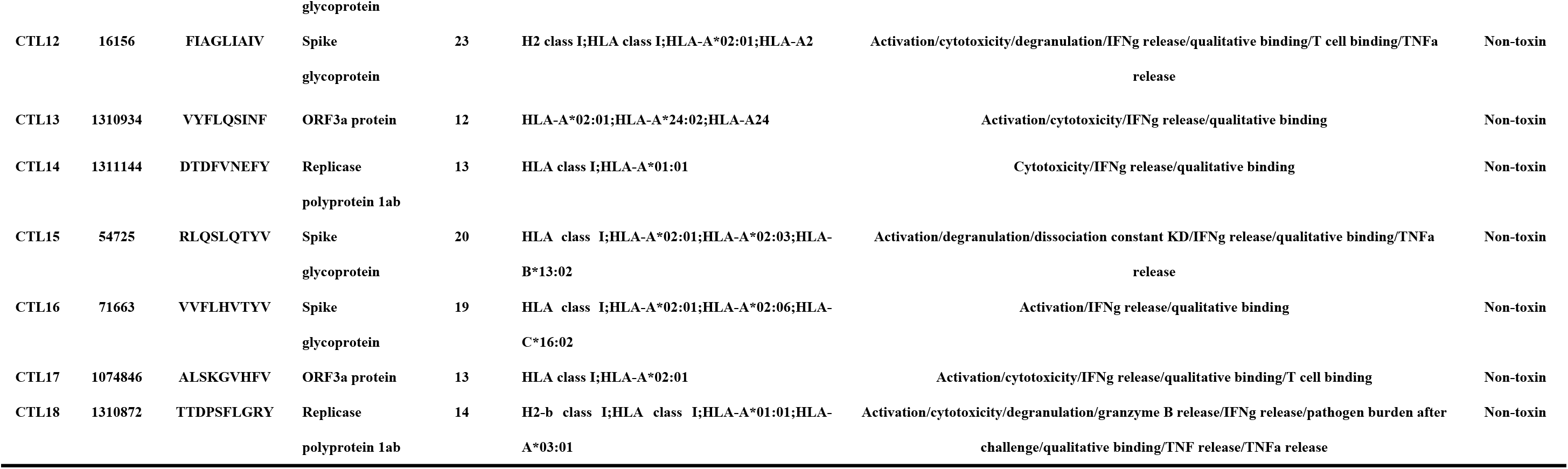
Cytotoxic T-lymphocyte specific epitope for the candidate pre-emptive Pan-Coronavirus Vaccine as well as toxin prediction for each epitope.

### The cross-reactivity of candidate CTL epitopes to its corresponding variants were determined through a structure-based immunogenic footprints analysis approach

The potential for immune evasion exists among the variants of candidate CTL epitopes, which reduces the effectiveness of the vaccine, while some CTL epitopes have shown cross-reactivity with their corresponding variants. To determine the cross-reactivity of these candidate CTL epitopes targeted by memory CD8^+^ T cells from SARS-CoV-2 variants and the 33 human and animal coronaviruses, it is necessary to further understand T-cell-mediated adaptive immune responses and design broad-spectrum antiviral vaccines that define the scope of immune protection. In this study, we employed structural modeling in combination with free energy prediction and surface electrostatic potential analysis to investigate immunogenic footprints left by candidate epitopes, aiming to identify cross-reactive targets with SARS-CoV-2 epitopes and evaluate the cross-protection potential of these candidate CTL epitopes (Figure 2C). Specifically, we focused on HLA-A*02:01-restricted CTL epitopes, which are representative of more than 50% of the human population, in order to investigate shared immunogenic features between the targets of SARS-CoV-2 variants and other coronaviruses (Figure S1). We selected 10 highly conserved cross-reactive CD8^+^ T cell epitopes from SARS-CoV-2 (Table 3) based on their binding affinity with HLA-A*02:01 molecules. These epitopes were derived from the Nucleocapsid protein, Surface glycoprotein, ORF1ab, ORF3a, and Membrane glycoprotein. Corresponding epitopes from SARS-CoV-2 variants of VOC and VOI, and other 33 alpha- and beta-coronaviruses members, were modeled in tertiary structures of HLA-A*02:01 (pMHC), comparing their molecular surfaces. Our results revealed that most mutations led to changes in the topology and surface electrostatic potential of pMHC, indicating that mutations among CTL epitopes can potentially cause immune escape (data not shown). However, we observed similar electrostatic distributions in the surfaces of CTL8 (MEVTPSGTWL) and its corresponding variant (MEVTPSGTWF), which corresponds to SARS-CoV-2 variants of BQ.1, as well as CTL11 (NYNYLYRLF) and its corresponding variant (NNNYLYRLF), which corresponds to SARS-CoV-2 variants of B.1.640. Despite amino acid changes in these peptides, they suggest preserved T-cell receptor (TCR) recognition (Figure 2D). It emphasizes the cross-protection potential of these candidate epitopes, favoring an antigen processing pathway for their antigens against the variants of VOC and VOI. Additionally, we modeled the pMHC structure of candidate CTL epitopes from different alpha- and beta-coronavirus members. The cross-reactive peptides exhibited similar pMHC surfaces. Nine coronaviruses (HKU1, HKU24, HKU23, NL63, Riyadh/Ry141, MERS, MERS-related CoV, and HKU5) showed shared electrostatic distribution and topography despite sequence divergences with CTL15 (RLQSLQTYV). Four coronaviruses (Civet007, YNLF_31C, WIV16, BtCoV-Recombinant coronavirus) showed shared electrostatic distribution and topography despite sequence divergences with CTL10 (LLYDANYFL). Six coronaviruses (HKU4, MERS, HKU5, MERS-Related-CoV, HKU9, Rousrttus Bt-CoV-GCCDC1) showed shared electrostatic distribution and topography despite sequence divergences with CTL3 (SPRWYFYYL). One coronavirus (MP789) showed shared electrostatic distribution and topography despite sequence divergences with CTL1(YLQPRTFLL) (Figure 2E). Furthermore, the MM-GBSA binding energy analysis predicted that the virus strain’s putative epitopes strongly bind to HLA-A*02:01, suggesting their potential as actual triggers for cross-reactivity of candidate epitopes (Supplementary data 2). No substantial physicochemical alterations were observed that could abolish their TCR cross-recognition. The CD8^+^ T cell-mediated response not only targets highly conserved epitopes but also shows cross-recognition of mutation epitopes from SARS-CoV-2 variants and some zoonotic coronaviruses. This highlights the cross-protection potential of these candidate epitopes, favoring an antigen processing pathway for their antigens against zoonotic coronaviruses. Furthermore, we identified anchor residues of candidate antigenic epitopes through structural analyses and determined the impact of variants of concern on T cell epitope recognition. These anchor residues remain fully conserved in SARS-CoV-2 variants of VOCs and VOIs, as well as alpha- and beta-coronaviruses (Supplementary data 2).

Altogether, we identified a total of 12 B-cell epitopes, 16 Th-cell epitopes, and 18 CTL-cell epitopes that are highly conserved among newly discovered variants of SARS-CoV-2, including VOC and VOI, as well as 33 zoonotic coronaviruses. Many of these candidate epitopes are found within structurally constrained regions of both the structure and non-structure proteins of SARS-CoV-2. These regions are highly conserved and cannot mutate without compromising the functionality of SARS-CoV-2 (Figure 3A). Moreover, our analysis revealed that all the conserved epitopes either overlapped or were unique to different strains of zoonotic coronaviruses (Figure 3B-D). Additionally, our predictions indicate that these epitopes are non-toxic to the host, as depicted in Table 1-3. On the other hand, mutational inactivation of a given CD8^+^ T cell epitopes is expected to be compensated by the persistent response directed against unchanged co-existing CD8^+^ epitopes. Notably, certain mutations in epitopes may exhibit cross-reactivity, although some corresponding epitopes were distinct to candidate epitopes. This implies that the candidate antigenic epitopes have the potential to induce a broader range of immune protection. These conserved epitopes could increase the coverage of the vaccine as well as solve the problem of genetic variability, thereby initiating new efforts for vaccine development.

**Figure 3.**
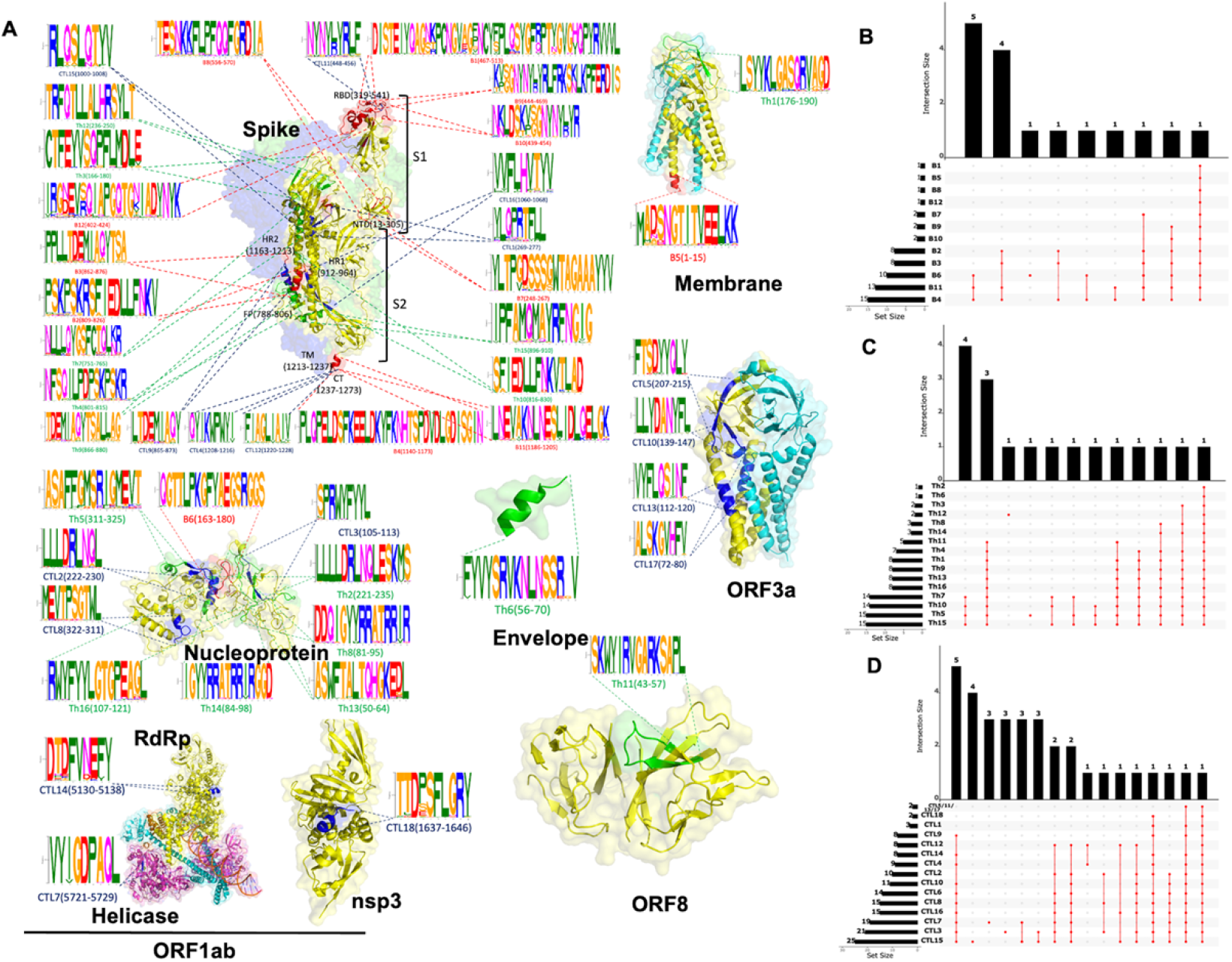
Candidate epitopes mapping to SARS-CoV-2 structural and non-structural proteins and the conserved relationship with different coronavirus strains. **(A)** Mapping of candidate epitopes to structures of surface spike glycoprotein (PDB ID: 6vsb, 3.46 Å), ORF1ab_nsp3 (PDB ID: 6w9c, 2.70 Å), ORF3a (PDB ID: 6xdc, 2.90 Å), ORF8a (PDB ID: 7jx6, 1.61 Å), Envelope protein (PDB ID: 7tv0, 2.60 Å), Membrane glycoprotein (PDB ID: 7vgr, 2.70 Å), Nucleocapsid (PDB ID: 8fd5, 4.57 Å) and ORF1ab_RdRp, Helicase (6xez, 3.50 Å). Every candidate epitope motif’s logo and the position of structure. The red, blue, and green colors represent B cell, CTL, and Th epitopes, respectively. **(B-D)** Conservativeness analysis of B, Th, CTL candidate epitopes in 33 zoonotic coronavirus strains. The left bar graph indicates the number of strains corresponding to fully conserved or cross-reactivity epitopes for each candidate epitope; the upper bar graph suggests the number of virus strains mapped to the same strain among different epitopes; and the middle-dotted line connecting graph refers to the number of strains corresponding to the intersecting viruses among different epitopes.

### The MHC polymorphism restricted of candidate T-cell epitopes, providing global population coverage

A given T-cell epitope can only elicit a response in individuals who express the corresponding specific MHC molecule. Therefore, selecting multiple epitopes with different HLA binding specificities can significantly enhance coverage of the patient population targeted by epitope-based vaccines or diagnostics. We have verified that all T-cell epitope are Pan-DR helper T cell epitopes and Pan-HLA-A CTL T cell epitopes. A total of CTL candidate epitopes that can be presented to CD8 T cells by 32 of the most prevalent HLA-A alleles. Additionally, we have identified Th candidate epitopes restricted by 50 of the most prevalent HLA-DRB / DQB / DQA / DPB alleles, as shown in Table 2 and Table 3. It is worth noting that these epitopes can cross-bind with high or intermediate affinity to the aforementioned HLA-A alleles. Notably, the HLA-A2-restricted candidate epitopes can also be cross-presented by mouse H-2K/Db molecules, which provide feasibility for our mouse-based animal model to validate vaccine immunogenicity. Population coverage analysis reveals that the selected T-cell epitopes provide coverage for 100% of the global population, as well as individual regions such as Northeast Asia, Europe, North America, Southeast Asia, East Africa, and China. East Asia, representing the main epidemic zone for SARS-CoV-2, has a coverage rate of 99% (Table S7). The allelic frequency data confirm the global distribution characteristics of the selected T-cell epitopes, making them suitable for designing candidate protein vaccines.

### Candidate vaccines have good immunogenicity and stability based immune-informatic and structure based Molecular dynamic simulation analysis

We utilized a combinatorial approach to design an all-in-one multi-epitope pre-emptive pan-coronavirus vaccine candidate. It involved incorporating highly conserved B- and T-cell epitopes from seven antigenic proteins of SARS-CoV-2 into the design. To overcome the limitation of CVB3 live attenuated viral vectors in terms of genome size, we constructed candidate vaccines by linking B-, Th-, and CTL-cell epitopes using flexible linkers KK, AAY, and GPGPG. These linkers were selected based on their ability to improve expression levels and biological activity (Figure 4A). The immunogenicity of the vaccines with different epitope combinations. Vaccines are constructed by combinations of selected optimal arrangements using immunogenicity as a parameter, named EB, ET, and EC, respectively. These vaccines consisted of 294, 288, and 254 amino acid residues, respectively, with molecular weights of 33 kDa, 32 kDa, and 26 kDa. AllerTOP v2.0 predictions indicated that all three vaccine candidates were non-allergenic. The predicted half-lifes within mammalian in vitro reticulocytes, yeast, and E. coli were provided in Table S8. According to Expasy ProtParam analysis, the vaccine proteins were defined as stable proteins with instability indices of 33.25, 39.28, and 21.92, while the aliphatic indices were 66.33, 80.94, and 66.73, respectively. Next, we predicted the tertiary structural modeling of the candidate vaccines using advanced ESM-fold and refined the models using the GalaxyRefine2 tool (Figure 4B).

**Figure 4.**
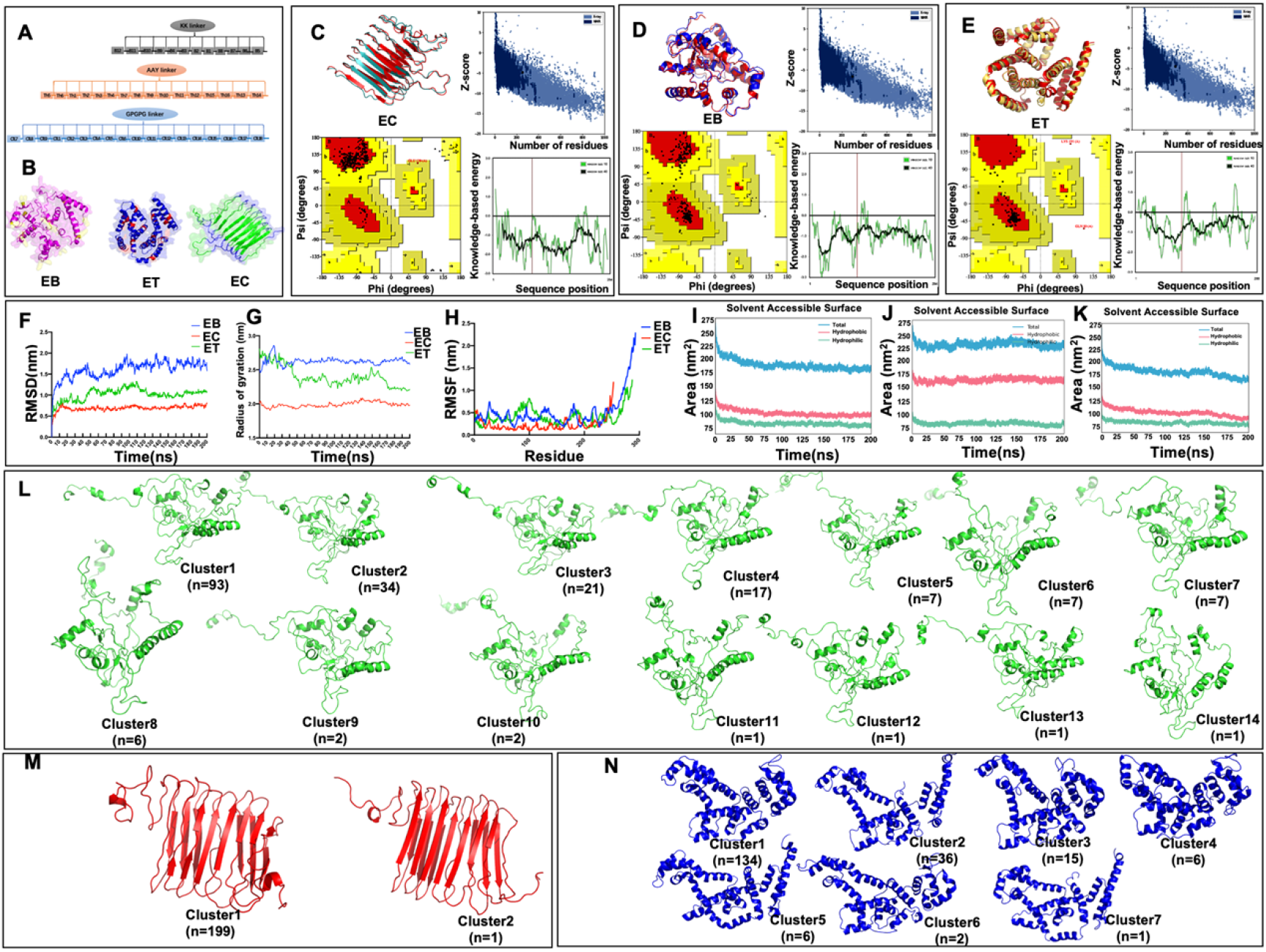
The candidate vaccines have good immunogenicity and stability by immune-informatics and molecular dynamic stimulation approaches analysis. **(A)** Schematic diagram of multi-epitope vaccine construction and **(B)** the model chosen through tertiary structure modeling for the multivalent epitopes subunit vaccine. Yellow, red and blue colors indicate KK, AAY, GPGPG linker, respectively. **(C-E)** Refinement and validation of tertiary structure models. Comparison between tertiary protein structures. The refined models are colored red; Ramachandran plot shows the presence of amino acid residues in favored, allowed and outlier regions; The Z-Scores of the best model in the ProSA-SEB plot were found to be -4.81, -4.46 and -6.58, respectively, which is within the range of conformational native proteins; The plot shows local model quality by plotting energy as a function of amino acid sequence position. **(F)** RMSD plot was obtained for the vaccine protein backbone after forming a complex with the EC, ET and EB proteins. The RMSD plot shows that it reaches equilibrium within 5–10 ns, with an RMSD value of 0.45 (nm); **(G)** RMSF, RMSF results showed the fluctuation of side chain residue of EB, ET, EC vaccine protein. **(H)** Radius of gyration (Rg) analysis. and **(I-K)** Solvent accessible surface (SAS) of EB, EC, ET analysis, respectively. **(L-N)** Structure conformation cluster analysis of EB, EC, ET vaccine proteins, respectively. The RMSD cutoff value is 0.5nm.

The best model was selected based on minimum galaxy energy. After refinement, the quality of the models were significantly improved, with more than 99% of residues falling within favorable regions (Table S9, Figure 4C-E). This refinement process was valuable in improving the quality of the structures obtained through ab initio model prediction. The modeled EB and ET structures, which maintained the minimum energy conformation, displayed a globular structural conformation primarily governed by α- helices and random coils. In contrast, the modeled EC structure exhibited a β-sheet repeat structural conformation. The GPGPG linker effectively linked CTL epitopes to form a stable structure, and the loop structure of linker helps the proteasome to process and present antigen epitopes. In conclusion, we designed an engineered vaccine that incorporated multiple B, CD4^+^, and CD8^+^ T-cell epitopes chimera of SARS-CoV-2. To assess B-cell induction, we predicted conformational B-cell epitopes using DisoTope 2.0. Among the three vaccines, EB possessed the highest number of conformational B-cell epitopes, followed by ET, while EC had the fewest. A total of 127 B-cell epitope residues were identified out of the 254 total residues of vaccine EC, 204 B-cell epitope residues were identified out of the 294 total residues of vaccine EB, and 9 B-cell epitope residues were identified out of the 288 total residues of vaccine ET (Figure S2 and Supplement data 3). These conformational B-cell epitopes corresponded to the exposed regions on the surface of the candidate vaccine proteins. The structural stability and flexible characteristics of protein vaccines play a crucial role in vaccine-induced host immune responses during antigen processing and presentation. Therefore, we conducted molecular dynamics simulations to evaluate the conformational stability of the candidate vaccine models. The analysis included the free energy landscape, as measured by RMSD and Rg. The RMSD results indicated significant structural changes in the three proteins at approximately 10 ns, which stabilized after 15 ns with minor fluctuations (Figure 4F). The EC protein remained stable throughout the 200 ns simulation, while the RMSD of EB protein exhibited more fluctuation. Overall, the RMSD values for all three vaccines fluctuated within the range of 5-20 Å, and the three vaccines designed were generally stable. The RMSF results indicated highly flexible regions at the C-terminal of all three proteins, with fluctuations between 5 and 25 Å. However, the other structural domains exhibited fluctuations within 10 Å. Solvent-accessible surface analysis revealed that vaccines EB and ET had the largest solvent-accessible surfaces at the 200 ns time scale (Figure 4G). The Rg analysis , which serves as a measure of the prime-weighted average radius of the system during simulation, showing differences in the compactness of the candidate antigens EB, EC, and ET over kinetic time scales, consistent with changes observed in RMSD (Figure 4H). Vaccine EB displayed sufficient flexibility and solvent-exposed characteristics to facilitate efficient presentation of B-cell antigenic epitopes (Figure 4I-K). Furthermore, conformational cluster analysis was done in RMSD cutoff value 0.5nm, with every 1ns trajectory Conformations. Result showed that most of the epitopes maintained their secondary structure in the three vaccines, although EB exhibited the largest number of conformational changes. Notably, the C-terminal domain displayed structural variability (Figure 4L-N). Overall, the MD simulation results confirmed that our designed vaccine meets the requirements of the principles of good antigen design.

### Attenuated CVB3 vectored intranasal pre-emptive pan-coronavirus vaccine induced System and mucosa immune responses by rCVB3-EPI and rCVB3-RBD-trimer in immunized mice

To guide our approach in developing a mucosal pre-emptive pan-vaccine, we constructed attenuated CVB3-based intranasal pre-emptive pan-coronavirus vaccines named rCVB3-EB, rCVB3-ET, and rCVB3-EC. Additionally, to characterize the antigenicity of the pan-coronavirus vaccines, we constructed an attenuated CVB3-based RBD trimer of SARS-CoV-2 (Wuhan-1) by the same strategy, ), referred to as rCV-RBD-trimer, as a positive control. Notably, to enhance the immunogenicity of RBD, we constructed a trimeric form by linking the Foldon domain to the C-terminus of RBD using a (GGGGS)_3_ linker (Figure 5AB). To improve the stability and expression efficiency of the designed pan-coronavirus vaccines in the murine expression system, we performed in silico cloning and obtained codon-optimized sequences for murine expression hosts. We tested the antigenicity and stability of candidate vaccine expression using Western Blot analysis, which confirmed the stable expression of RBD-trimer, EC, ET, and EB for at least 4 passages, along with their SARS-CoV-2 antigenicity (Figure 5C). Furthermore, we identified the plaque morphology of the recombinant virus-based vaccines and compared them to wild-type CVB3 using plaque assays. It was observed that rCVB3-RBD-trimer, rCVB3-ET, rCVB3-EC, and rCVB3-EB exhibited significantly smaller plaque morphologies compared to CVB3-WT, indicating potential attenuation of virulence in the candidate live virus vaccines (Figure 5D). In summary, we successfully constructed chimeric CVB3 vaccines with weak virulence and stable expression as candidates. Balb/c mice were used to evaluate the immunogenicity of candidate mucosa vaccine. A group of mice received rCVB3-RBD-trimer alone were employed as a positive control, while another group received rCVB3 alone were employed as a negative control. The timeline for the entire immunization and sample collection process is depicted in Figure 5E.

**Figure 5.**
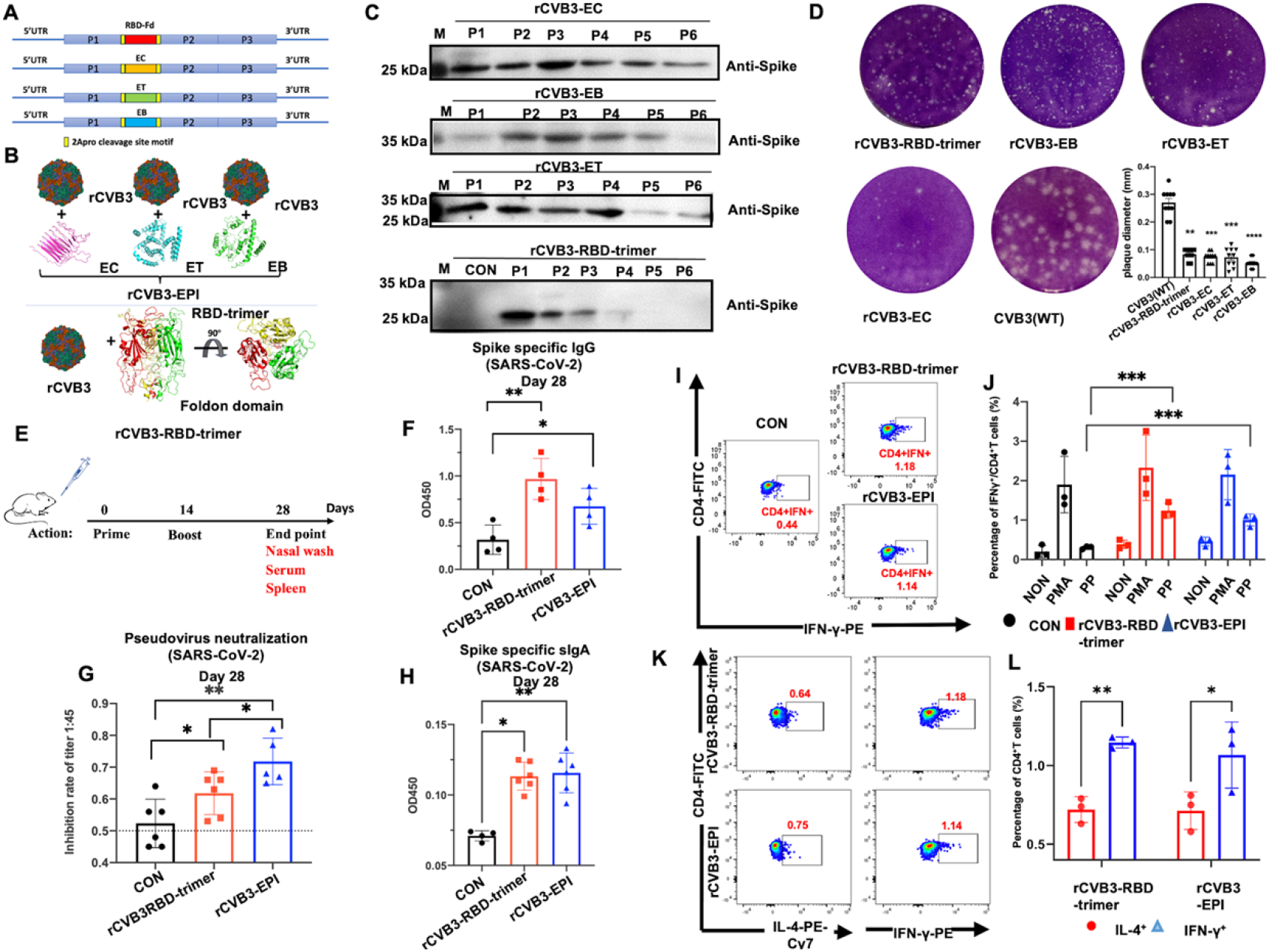
Attenuated CVB3 vectored intranasal pre-emptive pan-coronavirus vaccine induced system and mucosa immune responses by rCVB3-EPI and rCVB3-RBD-trimer in immunized mice. **(A)** Genome Schematic diagram of attenuated rCVB3 delivery vector based intranasal pre-emptive pan-coronavirus vaccines construction. **(B)** Cartoon models of rCVB3-EPI and rCVB3-RBD-trimer. **(C)** Western Blot analysis of the intranasal pre-emptive pan-coronavirus vaccines. Collection and analysis the protein in serial passages of the recombinant viruses (1-6 passages) from Vero culture cells. **(D)** Plaque morphology of viruses on Vero cells. For each recombinant virus, we randomly measured 10 Plaque diameter for statistical analysis. **(E)** Experimental schema of vaccination and sampling timeline. Nasal washes and serum were collected on day 28, n=6 /group. **(F)** SARS-CoV-2 S1 specific serum IgG antibodies levels on day 28, group rCVB3-RBD-trimer and rCVB3-EPI SARS-CoV-2 S1 specific serum IgG antibodies levels on day 28 compare to control group. IgG levels measured by ELISA and presented as OD450 nm values, n=4 / group. **(G)** The neutralizing activity of serum samples isolated from vaccinated mice (n =6 per group) against SARS-CoV-2 (Wuhan-Hu-1) pseudovirus. The vertical axis represents the pseudovirus inhibition rate of each immune serum at a serum dilution of 1:45. **(H)** SARS-CoV-2 S1-specific secretory IgA (sIgA) response in nasal lavage fluid of immunized mice was assessed on day 14 post booster immunization. n=6/group. **(L-J)** Percentages of IFN-γ^+^ CD3^+^ and CD4^+^ splenic T cells by FCM analysis on day 14 after the last immunization. The percentage of IFN-γ^+^ CD4^+^Th cells among from splenocytes was measured under non-stimulation (NON), PMA stimulation (PMA), and SARS-CoV-2 peptide pool (PP) stimulation, respectively, for both experimental group and vaccine group (n=3/ group). **(K-L)** The percentage of IL-4^+^ Th cells and IFN-γ^+^ Th cells among CD4^+^ T lymphocytes was determined under SARS-CoV-2 PP stimulation, respectively. Significance analysis was performed using a two-tailed t-test for unpaired samples. One-way analysis of variance (ANOVA) was also employed for significance analysis, data are represented as mean ± standard error of the mean (SEM) *, P<0.05; **, P<0.01; ***, P<0.001.

Comparatively, the rCVB3-RBD-trimer group and rCVB3-EPI group induced significantly specific IgG antibodies, while the rCVB3-RBD-trimer group induced significantly higher levels of specific IgG antibodies (Figure 5F). To quantify the neutralizing antibodies in mouse serum, we employed the SARS-CoV-2 (Wuhan-Hu-1) pseudovirus. Since some serum samples had neutralizing antibody titers exceeding the detection limit, we utilized the inhibition rate of each group’s serum against the SARS-CoV-2 (Wuhan-Hu-1) pseudovirus at a serum dilution of 1:45 to determine neutralizing antibody titers. The results showed that both rCVB3-EPI and rCVB3-RBD-trimer induced significantly higher levels of neutralizing antibodies compared to the rCVB3 control group, with statistical significance observed in both groups. These findings suggest that nasal immunization with both rCVB3-RBD-trimer and rCVB3-EPI effectively induces SARS-CoV-2 (Wuhan-Hu-1) specific neutralizing antibodies in mice and elicits humoral immune protection. Notably, rCVB3-EPI can induce stronger systemic SARS-CoV-2-specific immune protection through enhanced production of neutralizing antibodies compared to rCVB3-RBD-trimer, despite inducing lower levels of binding IgG (Figure 5G), which highlighted the advantage of epitopes-based vaccines for the enrichment of neutralizing antibody epitopes. Next, we quantified the SARS-CoV-2-specific sIgA in nasal wash samples from immunized mice. Compared to the rCVB3 control group, both the rCVB3-RBD-trimer and rCVB3-EPI groups exhibited significantly increased levels of SARS-CoV-2-specific sIgA on the 14th day after boost immunization (Figure 5H). These results indicate that both rCVB3-RBD-trimer and rCVB3-EPI significantly induce specific mucosal immunity. Due to MHC polymorphism and the need to simultaneously present epitopes recognized by CD8 T cell populations of different MHC allele restrictions (Table 2 and Table 3), we used flow cytometry to detect the levels of IFN-γ and IL-4 secreted by spleen-specific T lymphocytes on the 14th day after boost immunization. Spleen cells from each mouse in each group were divided into groups based on different stimulations: PMA stimulation (PMA), SARS-CoV-2 peptide pool stimulation (PP), and no stimulation (NON). Flow cytometry fluorescent antibody staining was performed after stimulation or no stimulation. The results revealed a significant increase in the number of IFN-γ-and IL-4-secreting lymphocytes in the rCVB3-RBD-trimer group compared to the other two groups during SARS-CoV-2 PP stimulation. However, the rCVB3-EPI group exhibited an upward trend in IFN-γ^+^ Th cells without statistical significance. These findings suggest that both rCVB3-RBD-trimer and rCVB3-EPI induce SARS-CoV-2 spike-specific Th1 lymphocytes (Figure 5I-J). It is important to note that the proportion of Th cells secreting IFN-γ was higher than those secreting IL-4 in both rCV-RBD-trimer and rCV-EPI groups (P<0.01), indicating the vaccine’s ability to promote Th1 T lymphocyte maturation and favor a Th1-type immune response. The enhancement may lead to antiviral effects through cytotoxic T lymphocytes (Figure 5K-L). Therefore, these results demonstrate that rCV-EPI can stimulate robust local mucosal immune responses, creating a protective environment in the respiratory tract. Overall, these findings support the notion that rCV-EPI can effectively elicit both humoral and T cell-mediated immune responses in both the systemic and mucosal immune systems.

## Discussion

The global spread and evolution of SARS-CoV-2, along with its zoonotic nature and potential for spillover, make it challenging to predict the strain responsible for the next coronavirus pandemic. Therefore, the development of a safe and effective preemptive pan-vaccine is crucial to protect against future COVID pandemics [6,7,10]. In this study, we have compiled immunodominant B, Th, and CTL epitopes from several studies in the IEDB database. Based on the evolutionary trajectory of the virus, we integrated immune-informatics and computational structural biology approaches to screen the both conserved and non-toxin candidate epitopes. Some of these epitopes have structural constraints that limit SARS-CoV-2 evolution and are potentially invariant CD4^+^ and CD8^+^ epitopes, making them excellent candidates for next-generation vaccines [11,12]. We have employed a combinatorial approach to design an all-in-one pan-coronavirus vaccine with immunogenicity and stability characteristics that promote protective immunity and prevent viral escape. Epitope-based vaccines offer significant advantages in terms of safety and effectiveness compared to empirical based traditional vaccines [13–17]. As our knowledge, while there have been other studies using similar strategies for multi-epitope vaccine design, ours is the only research work that has experimentally verified the reliability of the vaccine [11,12]. Our comprehensive approach underscores the adaptability of our pre-emptive multi-epitope pan-coronavirus vaccine strategy, which can be readily modified to target mutant strains as well as new zoonotic bat SL-CoV that may emerge in the future. This adaptability is expected to expedite the implementation of future pre-emptive multi-epitope vaccines, mitigating localized outbreaks and preventing them from transforming into global pandemics.

Vaccines that elicit durable mucosal immune responses in the respiratory tract hold significant potential in preventing virus infection, replication, shedding, and transmission [1]. Mucosal immunity to SARS-CoV-2 at the primary site of viral shedding could impede forward transmission [1]and lower the incidence of COVID-19 infection, thereby reducing the emergence of variants, disease surges, and associated morbidity. Currently, numerous mucosal vaccines targeting SARS-CoV-2 have been developed using various vaccine platforms (proteins, viral vectors, live attenuated viruses, DNA, RNA, and inactivated viruses) and different modes of delivery (nasal and oral droppers, sprays, inhalers, nebulized delivery, and oral tablets) [18,19]. The early-stage development of the attenuated vector CVB3 in the laboratory has demonstrated its ability to activate robust mucosal and systemic immune responses, leading to vertical maternal-infantile immune protection. It suggests that our candidate vaccine may play a role in protecting newborns from coronavirus infection, making it a potential platform for developing a SARS-CoV-2 mucosal vaccine. Intranasal immunization with rCVB3-RBD-trimer and rCV-EPI induces specific immune responses against SARS-CoV-2, including the production of specific IgG binding antibodies with functional serum-neutralizing capabilities. Notably, these vaccines also elicit respiratory SARS-CoV-2-specific sIgA, which is an important marker of intranasal immune response. During early-stage SARS-CoV-2 infection, before the appearance of IgG, sIgA plasmablasts with mucosal homing potential spread peripherally and secrete IgA antibodies, playing a crucial role in capturing and neutralizing the virus to prevent transmission [20,21]. CD8^+^ T cells play a crucial role in eliminating pathogenic infections. Different coronaviruses exhibit distinct patterns of reactivity towards various targets, resulting in amino acid sequence variations. Numerous variants can lead to immune escape, which cannot be explained solely through sequence analysis. Examination of structural data over epitope sequence analysis could explain how candidate CTL epitopes induced memory CD8^+^T cell may produce a heterologous immunity response in a global scale against emergent diseases such as COVID-19, mitigating its full lethal potential, and paves the way for the development of wide spectrum vaccine development [22]. Furthermore, both rCVB3-EPI and rCVB3-RBD-trimer can induce specific cellular Th immune responses when stimulated with spike protein antigen PPs, indicating that T cell epitopes can play a role, and there may be effector T cell epitopes present in the spike region that provide strong memory protection. The presence of more IFN-γ^+^ Th cells than IL-4^+^ Th cells suggests that the vaccine mediates a Th1-propensity type of response, contributing to its antiviral effect. While MHC polymorphisms were limited to the mouse model, some candidate Th and CTL epitopes have been confirmed to bind to MHC alleles in mice, supporting our findings. However, further investigation is needed to test the effects of T cells using human HLA transgenic mice or clinical trials. Population coverage analysis based on HLA allele polymorphisms demonstrates that our vaccine candidate is suitable for all populations worldwide. Both rCVB3-RBD-tirmer and rCVB3-EPI vaccines developed in this study successfully induce specific systemic and local mucosal immune responses in mice through nasal drip immunization.

Nonetheless, our study had several limitations. Firstly, in vivo studies have only been conducted in mice, with relatively small group sizes. Before advancing this vaccine to clinical trials, more animal models must be included, acknowledging the physiological differences between animals and humans. The local environment of the human nasal and respiratory tracts is more heterogeneous, and the immune system is more complex, requiring demonstration of actual immunity through clinical trials. Secondly, pre-existing CVB3 antibodies in the population may expedite the clearance of CVB3 vectors from the body, resulting in reduced replication of rCVB3-RBD-trimer and rCVB3-EPI, ultimately leading to decreased expression of RBD and epitope peptides. Thirdly, further research is necessary to determine the durability of the immune response, particularly in enhancing immunization effectiveness against heterologous viruses. Fourthly, the experiments did not evaluate the effectiveness of the vaccine in terms of cross-immunoprotection against various mutant strains. Finally, the attenuated enterovirus vector-based rCVB3-RBD-trimer and rCVB3-EPI may also present a risk of revertant mutations, requiring further investigation for clarification.

## Conclusion

Both rCVB3-RBD-trimer and rCVB3-EPI were both able to significantly induce SARS-CoV-2-s specific system immunity as well as mucosal immunity, suggesting that these vaccines can effectively impede viral invasion, lower the likelihood of severe acute respiratory disease, and mitigate virus transmission. Our research may offer some protection from future zoonotic coronavirus outbreaks, laying the foundation for the development of pre-emptive pan-coronavirus vaccines in future.

## Materials and Methods

### Sequence and structures data retrieval

A total of 33 representative alpha- and beta-coronavirus genome and protein sequences were obtained from the NCBI database, which include the sub-genus: Setracovirus, Duvinacovirus, Merbecovirus, Sarbecovirus, Hibecovirus, Nobecovirus, Enbecovirus) as well as species originating from humans, bats, camels, civet cats, pangolins, and mice. The accession numbers for these sequences can be found in Table S10. Additionally, a collection of 106 protein sequences representing different lineages of SARS-CoV-2 variants announced by the WHO as Variants of Concern (VOCs) and Variants of Interest (VOIs) was downloaded from the NCBI virus database (Table S11). Furthermore, X-ray crystallography structures of Surface glycoprotein (PDB ID: 6vsb, 3.46 Å), ORF1ab_nsp3 (PDB ID: 6w9c, 2.70 Å), ORF3a (PDB ID: 6xdc, 2.90 Å), ORF8a (PDB ID: 7jx6, 1.61 Å), Envelope protein (PDB ID: 7tv0, 2.60 Å), Membrane glycoprotein (PDB ID: 7vgr, 2.70 Å), Nucleocapsid (PDB ID: 8fd5, 4.57 Å), and ORF1ab_RdRp, Helicase (PDB ID: 6xez, 3.50 Å) from SARS-CoV-2 were collected from the RCSB PDB database.

### Sequence alignment and Phylogenetic tree construction

The MAFFT v7.313 software was utilized to align protein and genome sequences derived from humans, bats, pangolins, civets, and camels [23]. Consensus aligned protein sequences from each virus strain were subjected to sequence variation analysis. For genome comparison, the optimal site model was selected based on the lowest Bayesian Information Criterion (BIC) using the ModelFinder software’s IQ-tree model selection tool [24]. The SYM + R4 model was chosen for nucleotide substitution, along with auto thread and free rate heterogeneity + R. For the whole coronavirus genome sequences, the GTR + R model with auto thread 4 was determined as the best substitution for maximum likelihood (ML) calculation based on the lowest BIC score. Phylogenetic trees were constructed using the ML method in IQ-tree (version 1.6.8). The consistency of the topology for coronavirus sequences was assessed using a bootstrap method with 1000 replicates. The final phylogenetic tree was edited using Interactive Tree Of Life (iTOL) (https://itol.embl.de) [25].

### Epitopes prospections and conservative analysis

The Immune Epitope Database and Analysis Resource (IEDB) [26] was accessed to prospect and retrieve experimental SARS-CoV-2 CD8^+^ T cell, CD4^+^ T cell, and B-cell epitopes, establishing the SARS-CoV-2 epitopes landscape dataset. To ensure cross-clade immunity against variants undergoing diverse evolutionary pathways, candidate epitopes highly conserved between currently circulating VOCs, VOIs, human coronaviruses, and animal coronaviruses were screened using the following criteria: (1) Linear B-cell epitopes demonstrated potent neutralizing antibody induction; (2) Validation of all candidate epitopes employed a range of analytical strategies; (3) T-cell epitopes elicited distinct biological responses and exhibited remarkable conservation across different strains. To analyze putative epitopes from SARS-CoV-2 variants, 106 representative variants of SARS-CoV-2 VOCs and VOIs were aligned, and the epitope conservancy analysis tool was used to compute the degree of epitope conservancy within a given protein sequence of SARS-CoV-2, set at a 70% identity threshold to obtain the frequency of concerning mutations and conserved epitopes [27,28]. Additionally, the 33 representative sequences of alpha- and beta-coronaviruses were inspected for putative epitopes by screening their protein correspondences and searching for amino acid identity with immunogenic targets described for SARS-CoV-2. The degree of conservation evaluated the fraction of protein sequences containing regions conserved to epitopes. The NetMHCcons tool was employed to predict binding affinity in both discordant SARS-CoV-2 sequences and putative epitopes of other H-CoVs and SL-CoVs [29]. All candidate epitopes were tested for immunogenicity using IEDB[30], while toxicity was confirmed by ToxinPred [31].

### Viral immunogenic footprints and Population Coverage analysis

The panel of mutated amino acids reported in SARS-CoV-2 VOCs and VOIs variants were screened to evaluate their binding with the HLA-A*02:01 epitopes included in our analysis. Putative epitopes for alpha- and beta-coronaviruses, as well as SARS-CoV-2 variants of VOCs and VOIs, were modeled in HLA-A*02:01 to investigate their TCR interaction surfaces and identify immunogenic footprints and shared structural features. Customized pMHC models were constructed using reliable AlphaFold 2.3.2, incorporating the analyzed peptides anchored to HLA-A*02:01 [32]. PIPSA server was employed to calculate the electrostatic surfaces and topological conformation of the generated pMHCs, verifying the impact of amino acid changes on the overall electrostatic distribution in these pMHCs[33]. The PIPSA server application generated colors based on charge distribution, ranging from -5 (negative charges in red) to 5 (positive charges in blue), with neutral charges displayed as white. Population coverage calculation was performed using the Population Coverage software hosted on the IEDB platform to evaluate the distribution of screened CD8^+^ and CD4^+^ T cell epitopes in the global population, considering both HLA-I and HLA-II alleles [34].

### Multi-epitopes pre-emptive pan-coronavirus vaccines design

To address the low immunogenicity of single epitopes and increase protection coverage, a beading strategy was employed to link B-cell, Th-cell, and CTL epitopes with KK, AAY, and GPGPG linkers, respectively. As a positive control, the RBD was trimerized by incorporating the Foldon domain of T4 phage at the C-terminus of the RBD [35]. Vexigen2.0 was used to predict the immunogenicity of the designed vaccines. VaxiJen, a prediction algorithm for alignment dependence and immunogenicity, was adopted based on the self-cross covariance (ACC) of protein sequences transformed into vectors of equal length [36]. AllerTop server, which predicts allergenicity, determines allergenic potential based on the s ACC of protein sequences converted into equal-length uniform vectors. AllerTOP offers high sensitivity, up to 94%, compared to other methods [37]. Finally, ExPASy was employed to predict molecular weight, isoelectric point, half-life, stability, and other physical and chemical properties. After comparing and evaluating the immunogenicity and physicochemical characteristics of the multi-epitopes all-in-one vaccines, the best combination vaccine was selected and rename EB, ET and EC, representing the candidate vaccines.

### Candidate vaccine tertiary structure modelling and Conformational B-cell epitopes prediction

To model the tertiary structure of the candidate vaccine, we utilized ESMfold since there was no suitable template available for reference. ESMfold employs an AI large language model to accurately infer the structure from the master sequence [38]. GalaxyRefine2 was employed to refine the tertiary structure [39]. The quality of the tertiary structure model was assessed using the RAMPAGE online server and ProSA-web. ProSA-web calculated the interaction energy of each residue with the rest of the protein to determine if it meets certain energy criteria. An overall quality score was computed by ProSA, and if the score falls outside the characteristic range for native proteins, it suggests potential errors in the structure [40]. The Ramachandran plot predicted energetically allowed and disallowed dihedral psi (ψ) and phi (φ) angles of the amino acid residues based on the Van der Waals radius of side chain atoms [41]. Conformational B-cell epitopes of the vaccines were predicted using DiscoTope 2.0. This method calculates surface accessibility (estimated in terms of contact numbers) and a novel epitope propensity amino acid score. The final score is obtained by combining the propensity score of residues in spatial proximity with the contact numbers [42]. A default threshold of 3.7 was used, along with sensitivity and specificity values of 0.47 and 0.75, respectively. N and O glycosylation sites were screened using NetNGlyc 1.0 and NetOGlyc 4.0 prediction servers, respectively[43,44]. The structural figures were generated using the PyMOL Molecular Graphics System, version 2.3.0.

### Molecular dynamics simulations

Molecular dynamics simulations of the three candidate vaccines were performed using GROMACS 2023, with the amber03 force field selected. Periodic boundary conditions (PBC) were applied in all directions within a cubic box of size 10 Å. The system was solvated with the SPC/E water model. Charges generated for our system were neutralized using Na^+^ and Cl^−^ ions at a concentration of 0.15 mM. The initial equilibrium phase involved running an NVT ensemble with a constant temperature of 300 K maintained by a Berendsen thermostat. Equilibration steps included energy minimization, NVT, and NPT for 10,000 ps, 500 ps, and 1000 ps, respectively. MD production was performed for 200 ns with integration steps of 2 ps, under constant pressure and temperature conditions. The Root Mean Square Deviation (RMSD), Radius of Gyration (Rg) for the backbone, Root Mean Square Fluctuation (RMSF) for the side chain, and Solvent Accessible Surface (SAS) were determined. A series of trajectories were obtained by cluster analysis. Images were created using a custom in-house script for PyMOL 2.3.0.

### Cell and virus

The CVB3 (Nancy strain) was generously provided by Dr. Qihan Li (Institute of Medical Biology, Chinese Academy of Medical Science and Peking Union Medical College). The attenuated CVB3 strain (rCVB3) and pCVB3 cDNA used in the experiment were constructed and stored in our laboratory. Vero cells were obtained from the Cell Bank of the Chinese Academy of Sciences (Shanghai, China) and maintained in DMEM (8122622, Gibco, USA) supplemented with 10% fetal bovine serum (PF09666, Lonsera, Uruguay) and 1 × penicillin/streptomycin (15140-122, Gibco, USA) at 37 °C with 5% CO2. Viral stocks were prepared from Vero cells when the cytopathic effect (CPE) reached 95%. The viral supernatant from three freeze-thaw cycles was stored in aliquots at -80°C.

### Construction of recombinant CVB3 vectored rCVB3-ET, rCVB3-EC, rCVB3-EC and rCVB3-RBD-trimer

The cDNA coding sequences for ET, EB, and EC were synthesized by GenScrip Co., Ltd. (Nanjing, China). Based on the cDNA clone of pCVB3, the genes encoding RBD-trimer, EC, ET, and EB were connected to the pCVB3 vector using Cla I and Afe I restriction enzyme sites, resulting in the creation of pCVB3-RBD-trimer, pCVB3-EC, pCVB3-ET, and pCVB3-EB cDNA clones. The accuracy of the final clones was confirmed by sequencing. Primers used in the experiment are listed in table S12. Vero cells were transfected with pCVB3-RBD-trimer, pCVB3-EC, pCVB3-ET, and pCVB3-EB using the Lipofectamine 3000 system (2533476, Invitrogen, USA). Three days post-transfection, recombinant viruses (rCVB3-RBD-trimer, rCVB3-EC, rCVB3-ET, and rCVB3-EB) were harvested. The titer of TCID50 was determined using the Reed-Muench method and plaque assay. Viral titration and plaque assay were performed as previously described [9]. Plaques were visualized by fixing the cultures with 10% formalin for 30 min and staining with 1% crystal violet.

### Western Blot

The expression of the designed candidate vaccines using the recombinant CVB3 vector was confirmed by Western Blot assays. For each passage, Vero cells were infected with 100 μl of rCVB3-RBD-trimer, rCVB3-EC, rCVB3-ET, and rCVB3-EB, respectively, and incubated at 37°C until 90% CPE was observed. Cells were harvested and lysed on ice for 5 min using ristocetin-induced platelet aggregation (RIPA) buffer (P10013B, Beyotime, China), and the proteins were quantified using the BCA method. Protein samples were loaded onto a 10% SDS-PAGE gel, analyzed by Western Blot, and incubated with the SARS-CoV-2 Spike Antibody Mouse Mab (40592-MM117, SinoBiological, China) and HRP-conjugated goat anti-mouse IgG (ab6789, abcam, UK). Detection was carried out using an Ultra High Sensitivity ECL Kit (HY-K1005, MCE, USA) and the relative optical density of protein bands was quantified using Quantity One Software from Bio-Rad (Hercules, CA, USA).

### Immunization of mice

Balb/c mice were obtained from Beijing Vital River Laboratory Animal Technology Co., Ltd. (Beijing, China). All experiments were conducted in accordance with the National Institutes of Health Guidelines for the Care and Use of Laboratory Animals and approved by the Animal Care and Use Research Ethical Standards of Shantou University Medical College (approval grant no SUMC2020-136). The mice were housed at a temperature of 21±2°C, with a humidity range of 30%-70%, and 12h light/12h dark cycles. Food and water were provided *ad libitum*. Groups of age-matched Balb/c mice (n=15/group) at six weeks old were nasally immunized with a 1:1:1 mixture of 2×10^6^ PFU/ml rCVB3-EB, rCVB3-ET, and rCVB3-EC (referred to as rCVB3-EPI) on days 0 and 14. Negative control mice received 60 μl of 2×10^6^ PFU/mL rCVB3, while positive control mice received 60 μl of 2×10^6^ PFU/mL rCVB3-RBD-trimer. Prior to each immunization and on the 14th day after vaccination, nasal washes and serum samples were collected after anesthetizing the mice with isoflurane (R510-22, RWD Life Science, China) (induction: 5%; maintenance: 2.5%).

### Splenocytes isolation and Flow cytometry analysis

T cells were analyzed using flow cytometry as previously described [45]. Balb/c mouse splenocytes were harvested two weeks after the second immunization and suspended in 2 mL of cold phosphate buffered saline (PBS). The spleens were finely minced (approximately 0.2 cm^2^) and crushed using a grinding rod, then passed through sequential screens with pore sizes of 100 μm and 70 μm (BD Biosciences, San Jose, CA). The cells were pelleted by centrifugation at 400 × g for 5 minutes at 4 °C. Red blood cells were lysed using RBC Lysis Buffer (00-4333-57, Invitrogen, USA) followed by another wash. The isolated splenocytes were diluted to 5 × 10^6^ / ml viable cells per ml in RPMI-1640 media (SH40007, Thermo Scientific, USA) supplemented with 10% (v/v) fetal bovine serum (FBS) and 2× antibiotic–antimycotic solution for 6 hours. Cell viability was assessed using trypan blue staining.Simultaneously, the spleen cell suspension from each mouse was evenly divided into three wells in a prepared 12-well plate, labeled as "stimulated-PMA," "stimulated-PP," and "end stimulated" (non-activated). A Cell Stimulation Cocktail (00-4970, Invitrogen, USA) at concentration 1× 2 μg/mL was added to the PMA stimulation group. SARS-CoV-2 Spike peptide Pool (PP003, SinoBiological, China) with concentration of 1 × 2 μg/mL was added to activate cells in the peptide pool (PP) stimulation group. the cells. After incubating at 37°C for 1 hour, the cells responded to the stimulation and released cytokines. Brefeldin A Solution (00-4506, Invitrogen, USA) was administered 4–6 hours before staining to block intracellular cytokine secretion. Cells were subsequently washed with 1× PBS and stained using the buffer (554656, BD Pharmingen, USA) for 15 minutes at 4°C. First, anti-CD16/CD32 (553141, BD Pharmingen, USA) was added (0.5 μg / 50 μL) to block non-specific binding, followed by BV510 anti-CD3e (563024, BD Pharmingen, USA), FITC anti-CD4 (557307, BD Pharmingen, USA), and APC anti-CD8a (561093, BD Pharmingen, USA). Next, cells were fixed and permeabilized using the Fixation/Permeabilization Kit (554714, BD Pharmingen, USA) to enable intracellular staining with PE Rat Anti-Mouse IFN-γ (554412, BD Pharmingen, USA) and PE-Cy™7 Rat Anti-Mouse IL-4 (560699, BD Pharmingen, USA). Flow cytometry data were acquired using an Aurora™ Flow Cytometer (Cytek, USA) and a total of 100,000 events were acquired and analyzed using FlowJo V10 software.

### Neutralization assays

A neutralization assay was conducted using pseudovirus coated with the S protein based on the backbone of the VSV G pseudotype virus. The sera obtained from different groups were used for measurement. Serum samples were inactivated at 56°C for 30 minutes. Pseudovirus-SARS-CoV-2 Fluc WT (DD1502-03, Vazyme, China) with a concentration of 1.3 × 10^4^ TCID50 / mL was preincubated with sera from immunized mice at continuous triple dilutions or with control sera. After a 90 min incubation at 37°C, the mixture was added to vero cells to detect viral infectivity. The medium was changed the following day, and 28 hours after infection, luciferase expression in infected cells was determined using the Bio-Lite Luciferase Assay system (DD1201, Vazyme, China) and an enzyme-labeled instrument (Epoch, Bio-TEK, USA). The neutralization titer is defined as the reciprocal of the serum dilution corresponding to a 50% inhibition rate on the pseudovirus by the serum. The effective neutralizing antibody titer for the vaccine was categorized as follows: < 40 for ineffective (no inhibitory effect), 40 - 256 for effective (inhibitory effect), and > 256 for highly effective.

### Enzyme-linked immunosorbent assay of SARS-CoV-2 IgG and sIgA

The detection of SARS-CoV-2 IgG and sIgA was carried out using an enzyme-linked immunosorbent assay (ELISA) kit (KIT007, Sinobiological, China). Mouse sera were serially diluted and added to the plates, which were then incubated at 37°C for 1 hour. Following this, the plates were washed three times with a wash buffer. Antibodies, specifically goat anti-mouse IgG (HRP)-conjugated antibody (ab6789, Abcam, UK) and goat anti-mouse sIgA HRP-conjugated antibody (ab97235, Abcam, UK), were diluted 1:10,000 in blocking solution and added to each well (100 μl/well). After incubating for 1 hour at room temperature, the plates were washed five times with the wash buffer and developed with 3,3′,5,5′-tetramethylbiphenyldiamine (TMB) for 25 minutes. The reactions were stopped by adding 50 μl/well of a 1.0 M H_2_SO_4_ stop solution. The absorbance was measured at 450 nm using a microplate reader. To determine the titer of RBD-specific antibodies induced by the candidate vaccine, serum samples were serially diluted and measured by titration.

### Statistical analyses

All statistical analyses were performed using GraphPad Prism 8.0 software. Comparisons among multiple groups were performed using a one-way ANOVA followed by LSD’s multiple comparison post hoc test. Comparisons between two groups were assessed using unpaired Student’s t-tests. For non-normally distributed data the nonparametric Kruskal–Wallis rank ANOVA with post hoc Dunn’s test was used. A significance level of P < 0.05 was considered statistically significant. The notation *P < 0.05, **P < 0.01, ***P < 0.001, ****P < 0.0001 was used to indicate the level of significance. "NS" denotes non-significant results.

## Supporting information

**Figure S1.** Population coverage calculation of HLA-A*02:01 alleles.

**Figure S2.** Conformational B-cell epitopes of the candidate vaccines EB, EC and ET. Conformational B-cell epitopes are colored red.

**Table S1.** The candidate immunodominant B-cell epitopes exhibited a high degree of conservation among SARS-CoV-2 VOCs and VOIs.

**Table S2.** The candidate immunodominant B-cell epitopes exhibited a high degree of conservation among zoonotic coronaviruses.

**Table S3.** The candidate immunodominant Th-cell epitopes exhibited a high degree of conservation among SARS-CoV-2 VOCs and VOIs.

**Table S4.** The candidate immunodominant Th-cell epitopes exhibited a high degree of conservation among zoonotic coronaviruses.

**Table S5.** The candidate immunodominant CTL-cell epitopes exhibited a high degree of conservation among SARS-CoV-2 VOCs and VOIs.

**Table S6.** The candidate immunodominant CTL-cell epitopes exhibited a high degree of conservation among zoonotic coronaviruses.

**Table S7.** Population coverage calculation of the helper T lymphocyte (HTL) epitope and cytotoxic T-lymphocyte (CTL) epitope specific allele.

**Table S8.** physicochemical properties prediction of candidate vaccines.

**Table S9.** Candidate vaccine tertiary structure modelling quality analysis and refining.

**Table S10.** List of sequences of 33 representative alpha- and beta-coronavirus genome and protein sequences used in this analysis.

**Table S11.** List of sequences of 106 representative SARS-CoV-2 variants of concern (VOCs) and variants of interest (VOIs) sequences used in this analysis.

**Table S12.** PCR Primers used in this study.

## Acknowledgments

This study was supported by Natural Science Foundation of Guangdong Province (2021A1515012470, 2022A1515010540), 2020 Li Ka Shing Foundation Cross-Disciplinary Research Grant (2020LKSFG01E), Shantou Science and Technology Bureau (Shanfuke (2021)88-STKJ2021197, Shanfuke (2020)53-51).

## Author contributions

G.W., R.L., and H.D. conceived this project. H.D., Y.L., G.L.,H.Z. and J.Z. conducted immune-informatic analyses. X.H. constructed the plasmids and produced recombinant viruses. X.H., H.W., S.W., H.Z. and J.Z. performed antigenicity and immunogenicity evaluate. H.D. and Y.L. managed the project. H.D., Y.L. analyzed the results and wrote the manuscript. G.W. and R.L. directed and supervised the project.

